# Conformational dynamics of α-synuclein during the interaction with phospholipid nanodiscs by Millisecond Hydrogen Deuterium Exchange Mass Spectrometry

**DOI:** 10.1101/2020.11.16.385187

**Authors:** Irina Oganesyan, Cristina Lento, Anurag Tandon, Derek J. Wilson

**Affiliations:** Department of Chemistry, York University, Toronto, M3J 1P3, Canada; Department of Medicine, University of Toronto, Toronto, M5S 1A1, Canada; Centre for Research in Mass Spectrometry, York University, Toronto, M3J 1P3, Canada

**Keywords:** α-synuclein, phospholipids nanodiscs, conformational dynamics, Parkinson’s disease, hydrogen deuterium exchange, mass spectrometry

## Abstract

Both normal and pathological functions of α-synuclein (αSN), an abundant protein in the central and peripheral nervous system, have been linked to its interaction with membrane lipid bilayers. The ability to characterize structural transitions of αSN upon membrane complexation will clarify molecular mechanisms associated with αSN-linked pathologies, including Parkinson’s disease (PD), Multiple Systems Atrophy and other synucleinopathies. In this work, Time-Resolved ElectroSpray Ionization Hydrogen/ Deuterium Exchange Mass Spectrometry (TRESI-HDX-MS) was employed to acquire a detailed picture of αSN’s conformational transitions as it undergoes complexation with nanodisc membrane mimics. Using this approach, αSN interactions with DMPC nanodiscs were shown to be rapid exchanging and to have a little impact on the αSN conformational ensemble. Interactions with nanodiscs containing lipids known to promote amyloidogenesis (*e.g*., POPG), on the other hand, were observed to induce substantial and specific changes in the αSN conformational ensemble. Ultimately, we identify a region corresponding residues 19-28 and 45-57 of the αSN sequence that is uniquely impacted by interactions with ‘amyloidogenic’ lipid membranes and may therefore play a critical role in pathogenic aggregation.

## Introduction

Alpha synuclein is a 140 amino acid (14.5 kDa) pre-synaptic protein (1, 2). The exact biological function of αSN remains unknown (3), but it is now widely accepted that it plays a role in cell signalling processes (4, 5) and works as a molecular chaperone for the SNARE complex for compartmentalization, storage, and recycling of neurotransmitters (6–12). Still, αSN is more known for its misfunction, as misfolding of this protein initiates the formation of aggregation-prone intermediates that are the main constituents in protein aggregates like Lewy bodies (LB) and Lewy neurites (LN); the hallmarks of Parkinson’s and some other neurodegenerative diseases (11–17). Although αSN falls into the category of intrinsically disordered proteins (IDPs) and has been shown to be deficient of significant amounts of a secondary structure under physiological conditions (14, 20), it can exhibit a variety of conformational ensembles in response to environmental conditions and/or ligand binding, including α-helical or β-structured monomers, aggregates, oligomers, spheroids, fibrils among other configurations (21, 22).

Conformational changes of αSN upon interaction with vesicle’s phospholipid membranes is suspected to play a central role in amyloidogenesis and neuronal cell toxicity during Parkinson’s disease (23–29). While free αSN is largely disordered, membrane interactions drive the conformational ensemble to favor helical configurations though a series of intermediates (30–35). These intermediates represent a gateway to ‘off pathway’ conformations that are prone to form β-sheet rich ‘stacked’ protofilaments, which ultimately leads to the formation of amyloid aggregates (36–43).

Several models have been proposed to rationalize the structural behaviour of αSN when interacting with synaptic vesicles (10, 44, 45). A consolidated model was introduced by Dikiy and Eliezer (36), which can be summarized as follows: The αSN conformational ensemble is in equilibrium between an intrinsically disordered ‘free’ state and a vesicle-bound state, corresponding to a partially α-helical monomer, in neurons. The α-helical form can interconvert from an ‘extended helix’ where N-terminus (residue 1 to 60) and non-amyloid-beta component (NAC) (residues 61-95) form one helix that interacts with the vesicle’s bilayer to a ‘broken-helix’ where the long helix is broken at around position 45, which is thought to facilitate docking of αSN-loaded vesicles for exocytosis. The process of interconversion between ‘extended’ to ‘broken-helix’ state is proposed to allow for the formation of an off-pathway conformational intermediates that are prone to aggregation due to the exposure of the hydrophobic NAC region, which templates amyloid aggregation of disordered αSN in the cytosol (36).

While these membrane-models provide a detailed account of αSN dynamics in early amyloidogenesis, they are often supported by limited biophysical data. This is because the conformational dynamics of αSN’s folding and misfolding pathways in the presence of phospholipid bilayers presents a nmajor challenge for classical high-resolution structural techniques such as X-ray crystallography and structural NMR, which are not exceptionally well suited to the analysis of IDPs, even without the complicating factor of membranes (46–51).

In this work, we characterize the conformational ensemble of αSN in the presence and absence of various model membrane nanodiscs (52–58) using TRESI-HDX-MS (59). TRESI-HDX-MS is similar to classical ‘bottom up’ hydrogen deuterium exchange experiments but uses millisecond – low second labeling times. This approach allows for the characterization of weakly ordered conformational ensembles, including IDPs (38, 60, 61). It has been applied in a wide range of systems involving rapid conformational changes including protein-RNA/DNA (62, 63) and lipid (64) interaction, enzymatic transition states (65), intrinsically disordered protein folding/misfolding intermediates (66), and antibody-antigen interactions (67, 68).

In the TRESI-HDX setup used in the current study (Figure 1), millisecond hydrogen deuterium exchange reaction times are made possible by a concentric capillary rapid mixer labelled as ‘HDX reaction chamber’ in Figure 1 (69–72). The microfluidic device that follows incorporates all of the sample handling processes required for a ‘bottom up’ HDX workflow, including acidification of the sample (to quench exchange), proteolysis using an acid protease (usually pepsin) and electrospray ionization. Using this approach, we are able to acquire unique insights into the shifts in conformational bias that occur when αSN interacts with model membranes, including nanodiscs consisting of 1,2-dimyristoyl-sn-glycero-3-phosphocholine (DMPC) and 1-palmitoyl-2-oleoyl-sn-glycero-3-phospho-(1’-rac-glycerol) (sodium salt) (POPG), and mixed systems. Our aim is to define which conformational biases are associated with pathogenesis in α-synucleinopathies.

**Figure 1.**
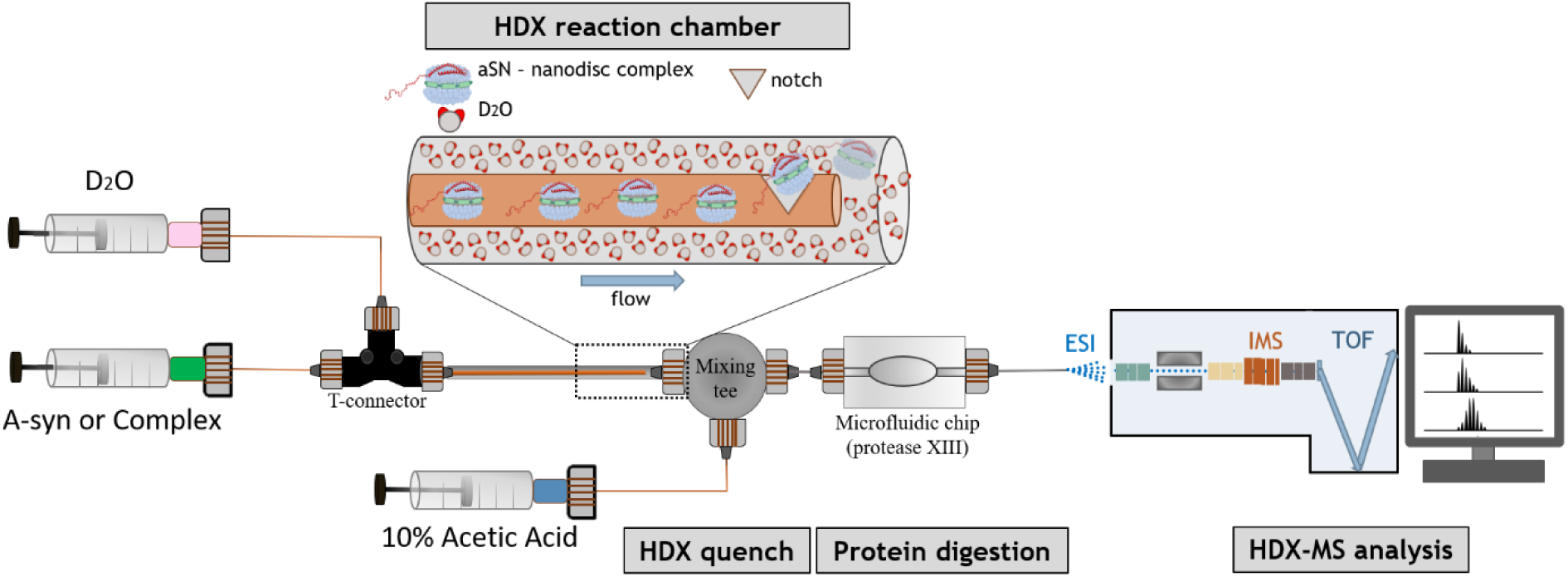
Experimental setup for the TRESI-HDX-MS workflow. Two syringes filled with D_2_O and premade αSN-nanodisc complex were hooked up to a T-connector by glass capillaries. The third end of the T-connector was attached to a metal capillary (bigger in diameter than the glass one). The complex’s glass capillary was threaded trough the metal capillary; D_2_O directly flowed inside the metal capillary. Complex and D_2_O met and mixed at the notch, which could be pulled back for varying volumes of the HDX reaction. This time-resolved mixer was connected to a mixing tee (grey circle), where the acid line was quenching the HDX reaction. Quenched αSN-nanodisc complex continued flowing through the microfluidic chip filled with protease XIII leading to protein digestion. Resulted peptides were introduced to nano-ESI on Synapt G2-S with IMS function on. HDX-MS analysis was performed using MassSpec Studio 1.0 Software.

## Results

### Protein Purification and Native ESI-MS

For the two main proteins required in the study, αSN and Membrane Scaffold Protein (MSP)1D1 ΔH5 (6 nm nanodisc belt), purity was assessed by SDS-PAGE (Figure S1) and ESI-MS (Figure 2). Gel fractions corresponding to 200 and 500 mM NaCl of anion exchange for αSN gave a clean band ∼15 kDa, and for MSP1D1 ΔH5 fractions (E5&E6) after Ni^2+^ affinity purification displayed strong band at ∼22 kDa. Both bands corresponded to the calculated and reported masses of the proteins (57, 73). The corresponding fractions were buffer exchanged *via* overnight dialysis against 0.1 M ammonium acetate pH 7.4 and analyzed *via* ESI-MS.

**Figure 2.**
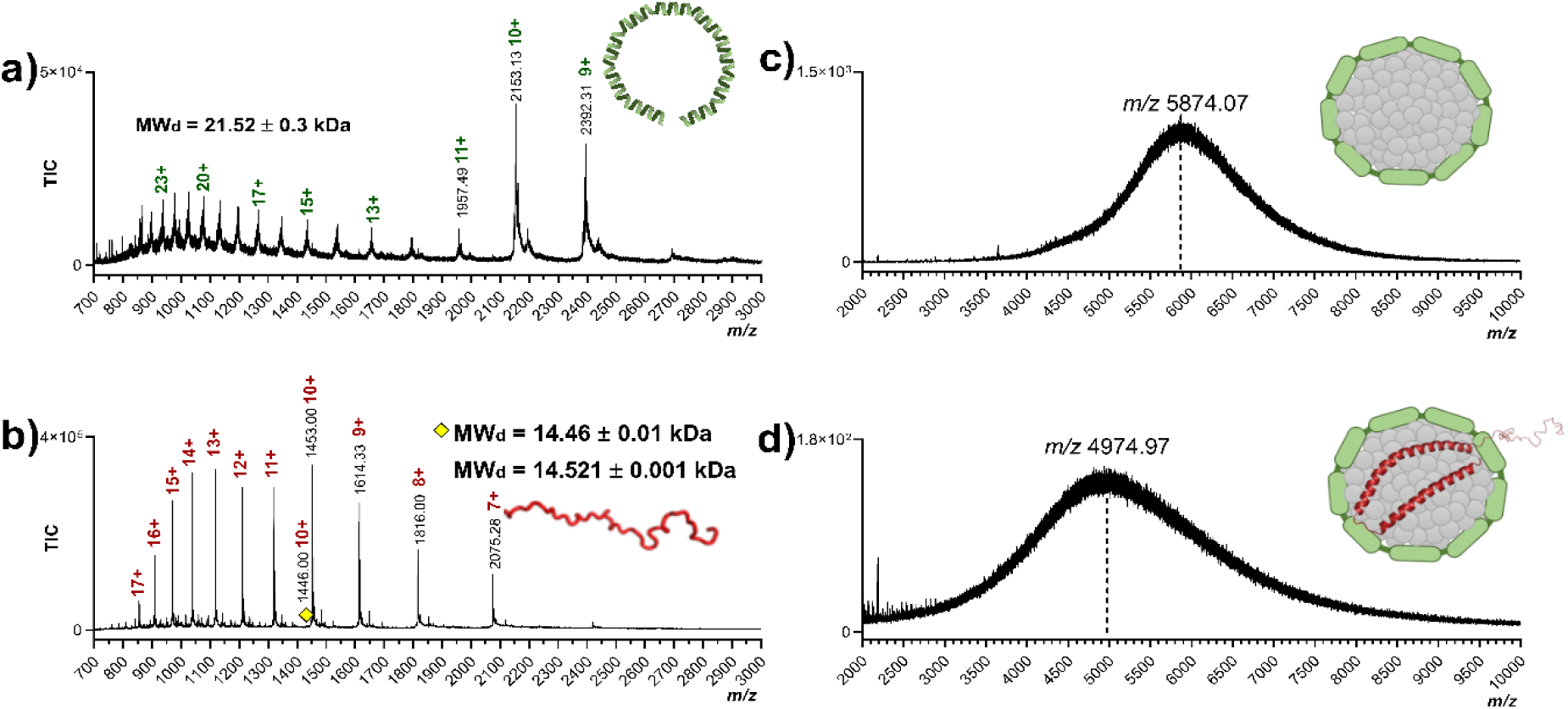
Native ESI mass spectra: **a**) MSP1D1 ΔH5 belt protein after expression/purification, **b)** αSN after expression/purification, **c)** MSP1D1 ΔH5 nanodisc after SEC separation, **d)** αSN/DMPC (MSP1D1 ΔH5) nanodisc complex after SEC separation. αSN and MSP1D1 ΔH5 proteins were diluted to 2.5 µM in 100 mM ammonium acetate pH 7.4 and infused into the Synapt G2-S mass spectrometer at a flow rate of 10 µL/min. Mass spectra were visualized by Prism 8.0.1, and deconvoluted using both UniDec and ESIProt software.

ESI-MS spectra of intact/undigested αSN and MSP1D1 ΔH5 were obtained in order to confirm the protein identity by mass and check for any impurities or degradation products. Figure 2(a) and (b) show the native MS spectra for MSP1D1 ΔH5 and αSN, respectively, along with the deconvoluted mass generated both by ESIProt (74) and additionally checked by UniDec software (75). αSN’s spectra exhibited a broad envelope of charge states in the lower *m/z* range, which is consistent with the intrinsically disordered state of the protein (Figure 2b) and previous ESI-based studies on αSN (76–78). The deconvoluted mass of the base peak of the 10+ charge state (yellow rhombus) was calculated to be 14.46 ± 0.01 kDa which matched the theoretical mass of an unfolded monomer (14.46 kDa) precisely (79). The most intense *m/z* peaks corresponded to Na^+^+ K^+^ (∼62 Da) adducts of the base peak, likely due to carryover from anion exchange purification.

The native MS spectrum for MSP1D1 ΔH5 exhibited two peak distributions – a broad envelope of charge states in the lower *m/z* range and a tight distribution in the higher *m/z* range, corresponding to a folded/unfolded equilibrium upon transfer to the gas phase (Figure 2a). The deconvoluted mass of MSP1D1 ΔH5 was 21.521 ± 0.034 kDa, which was in agreement with the calculated mass by sequence (including the His-tag and TEV site) of 21.552 kDa (57).

### Analysis of Nanodiscs and Nanodisc-αSN complexes using NMR, Native ESI MS and Size Exclusion Chromatography

Formation of nanodiscs was monitored by ^31^P NMR (Figure S2) and ESI-MS. In the case of NMR, intact nanodiscs were identified *via* a characteristic sharp peak at 0ppm (80–82). By ESI-MS, nanodiscs generate a broad ‘hump-like’ peak resulting from incomplete desolvation and polydispersity in the number of lipids contained in individual nanodiscs (Figure 2c) (83–85). Together, these data unambiguously demonstrate successful generation and purification of MSP1D1 ΔH5 nanodiscs.

Complexation of αSN with nanodiscs was explored by Size Exclusion Chromatography (SEC) and native mass spectrometry. Figure 3(a) illustrates the UV_280_ overlaid traces of a DMPC free nanodisc (blue) and free αSN (yellow) along with the SDS-PAGE bands corresponding to eluted protein(s) in specified fractions. The pure DMPC nanodisc chromatogram exhibited a broad peak from fraction B10 to B4 with a small shoulder around B11 to B10, both associated with the free nanodisc based on the SDS-PAGE bands of MSP1D1 ΔH5 and ESI-MS (86). A second smaller peak from C2 to C4 corresponded to unbound lipids and peptides (Figure 3a). Free αSN (orange trace), gave a first peak SEC around elutions B5 to B3 under our conditions, which was also verified by the SDS-PAGE and ESI-MS (Figure 3a). This relatively rapid elution for a small protein is consistent with the lack of compact structure of αSN. The second intense peak in the free αSN chromatogram (elutions B1 to C2) corresponded to a mixture of intact protein, peptides, and some small molecule impurities that were detected by ESI-MS.

**Figure 3.**
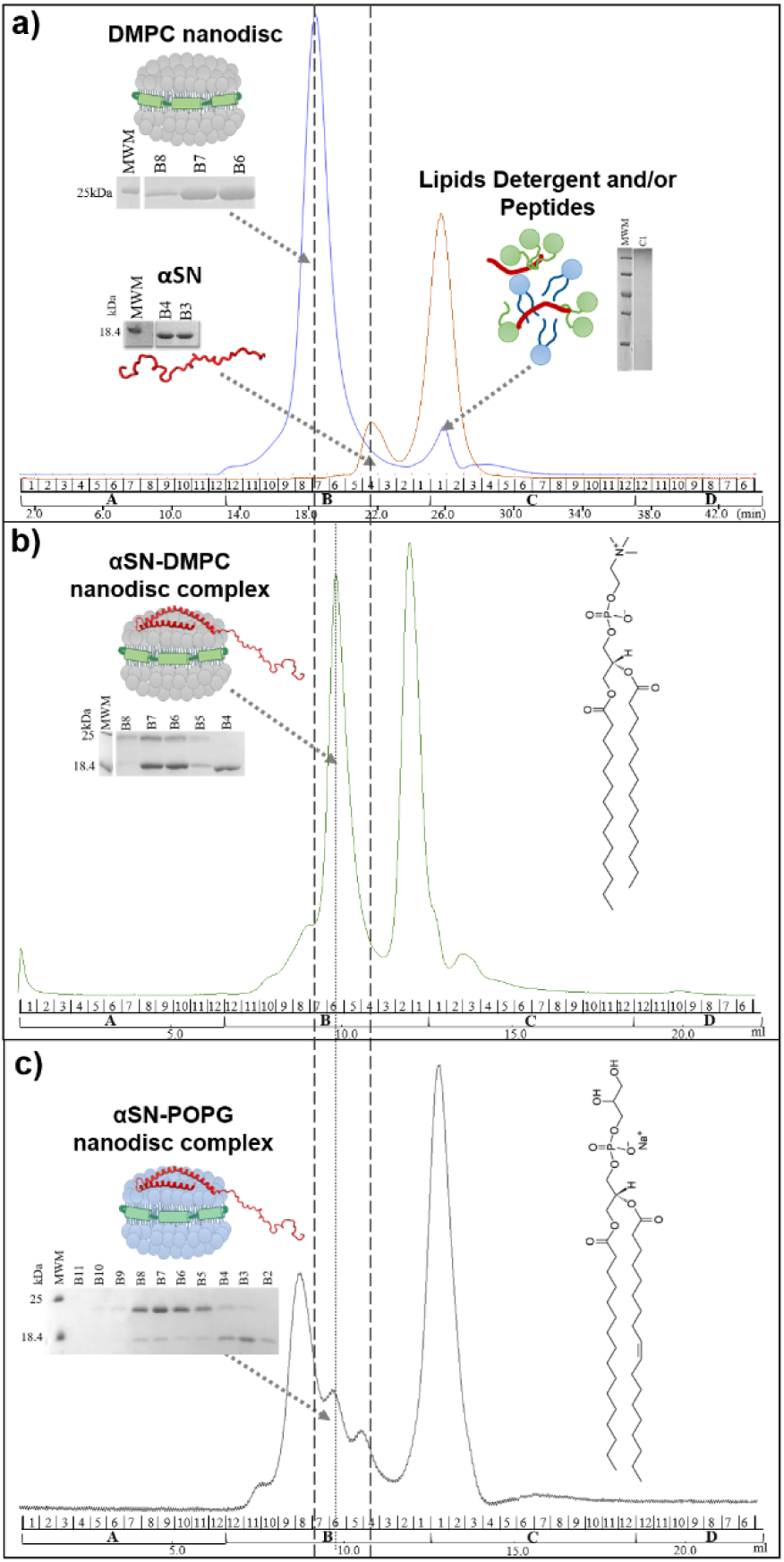
Size exclusion chromatograms and corresponding SDS-PAGE protein bands of samples: **a)** DMPC nanodisc complex (UV_280_ trace in blue) along with free αSN (yellow), **b)** αSN/DMPC nanodisc complex (green UV_280_ trace), and **c)** αSN/POPG nanodisc complex (grey UV_280_ trace). Samples were run on Tosoh Biosciences TSKGEL BioAssist G3SWXL 7.8 × 300 mm (5 μm, 250 Å) column with 0.1 M NH_4_OAc pH 7.0 at 0.5 mL/min. SDS-PAGE was performed using molecular weight marker from ThermoFisher (#PI-26616). Chemical formulas of DMPC and POPG finalize figure b) & c).

The αSN/DMPC nanodisc complex (green UV_280_ trace in Figure 3b) showed a profile consisting of two main peaks: B7-B5, and B2-C1, with the latter peak corresponding to a mixture of contaminants. SDS-PAGE bands for elutions B7 - B5 had indicated two bands matching MSP1D1 ΔH5 belt protein and αSN indicating the presence of a mixture of both nanodisc and αSN (Figure 3b). However, the peak sits between the ‘pure nanodisc’ and ‘pure αSN’ peaks shown in Figure 3(a), having a slightly longer retention time than the free nanodiscs and a slightly shorter retention time than free αSN. This is indicative of an interaction between DMPC nanodisc and αSN, but a ‘high turnover’ one where the exchange between ‘on’ and ‘off’ states is sufficiently fast to cause an averaging of the peak position in SEC (87, 88). ESI-MS data also provide additional evidence of an αSN/DMPC nanodisc interaction (Figure 2d), corresponding to a significant shift in the unresolved ‘hump’ to lower *m/z*. This can be attributed to the incorporation of additional positive charge carried by aSN into the complex (Figure 2d). However, the ESI process is known to generate both false positives and false negatives in protein interactions and therefore cannot in itself be taken as being conclusive. Nonetheless, the SEC and SDS-PAGE and ESI-MS results together provide strong evidence for rapid turnover αSN/DMPC nanodisc complexation.

In contrast to DMPC nanodiscs, the αSN/POPG nanodisc SEC profile exhibited a partially-resolved peak for the complex, corresponding to elutions B8 – B5 (Figure 3c). This peak was flanked to the left by a peak corresponding to the free nanodisc and to the right by a peak corresponding to free alpha synuclein, as determined by ESI-MS and SDS-PAGE. The occurrence of a distinct peak corresponding to the complex is suggestive of a ‘low turnover’ complex, where exchange between free and bound states is slower than (or on a similar timescale to) the SEC experiment.

### Deuterium uptake analysis of αSN/DMPC nanodisc, αSN/POPG:DMPC (1:1) nanodisc and αSN/POPG nanodisc complexes compared to the free αSN

To shed more light on how complexation influences the αSN conformational ensemble, differential TRESI-HDX experiments were carried out. In this approach, deuterium uptake in free αSN is subtracted from αSN deuterium uptake in the presence of nanodiscs. The result is ultimately a set of ‘percent deuterium uptake difference’ values which highlight changes in the conformational ensemble (measured as increased or decreased hydrogen bonding and/or solvent access in backbone amide hydrogens) in different regions of the protein upon complexation (Figure 4).

**Figure 4.**
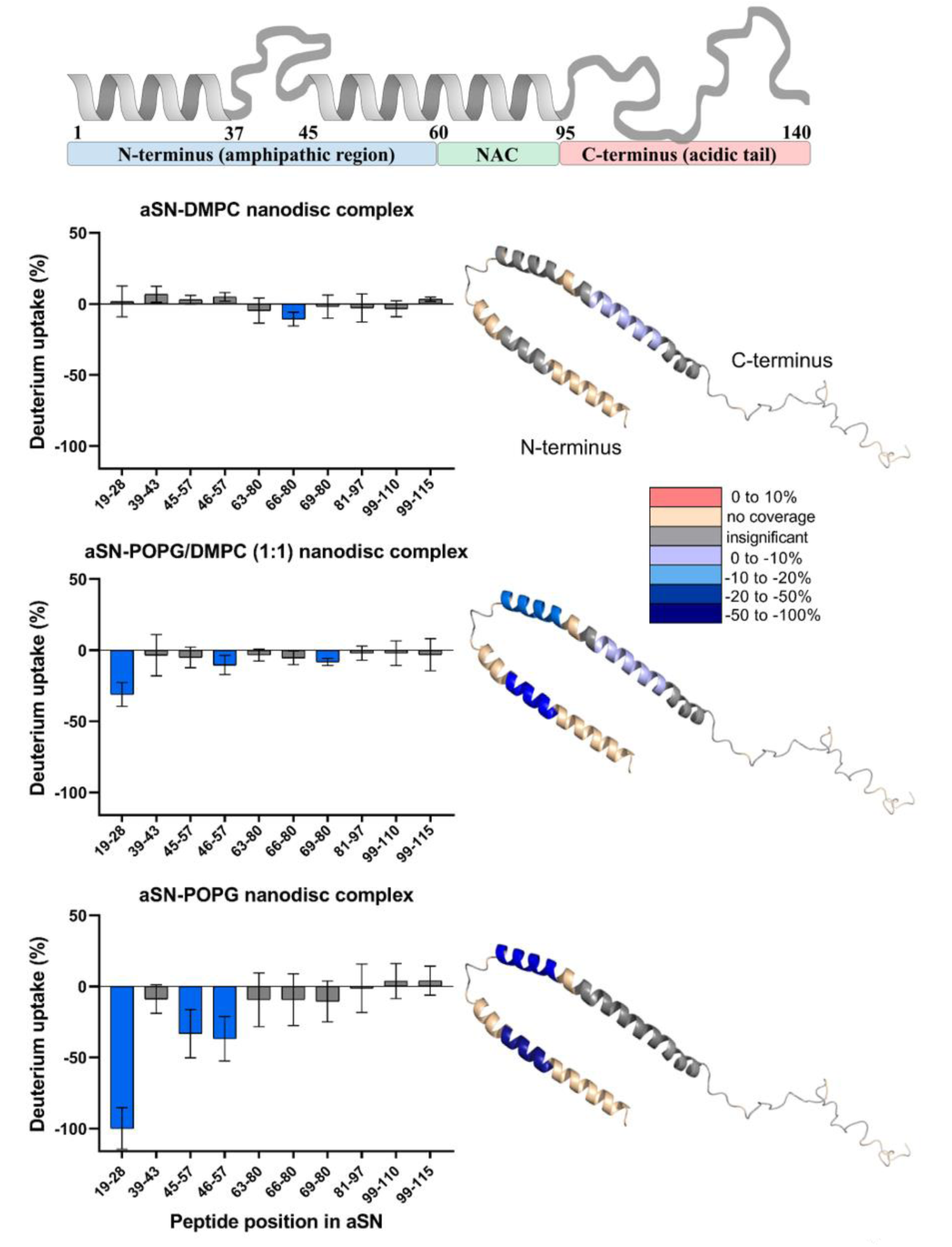
Sum of the all time points of TRESI-HDX difference measurements across 3 technical replicates comparing free αSN to αSN/DMPC nanodisc complex, αSN/POPG:DMPC (1:1) nanodisc complex, and αSN/POPG nanodisc complex (from top to bottom). For the graphs, positive (red) bars indicate increased deuterium uptake relative to the free protein and negative (blue) bars denote decreased deuterium uptake relative to the free protein. Grey bars represent peptides that do not have significant changes in deuterium uptake (> 1σ). To the right of each bar graph, deuterium uptake differences are mapped onto the corresponding crystal structure of αSN (PBD ID: 1XQ8). Gold region correspond to unreported coverage, gray to a non-significant differences, and blue and to red to a specified degree of increase or decrease in deuterium uptake.

In Figure 4, the differences in deuterium uptake were summed across six HDX labeling time points (0.12 sec, 0.34 sec, 0.66 sec, 1.32 sec, 2.2 sec and 3.3 sec), and averaged across three technical replicates for each timepoint. To provide a better view of the relative magnitudes of the observed differences between systems, all data were normalized to the highest observed difference (corresponding to segment 19 – 28 in the αSN/POPG system). Differences in deuterium uptake were mapped onto the corresponding crystal structure of αSN (PBD ID: 1XQ8) for easier visualization (35).

Overall, the αSN/DMPC nanodisc complex exhibited the lowest differences in deuterium uptake, with the majority of segments showing non- or barely significant changes (Figure 4, top). This profile is partially consistent with the SEC results described earlier, in the sense that the HDX signal for a high-turnover complex could strongly weighted toward the ‘off’ (*i.e*., ‘free’) state, and would therefore show limited or no difference compared to the ‘free’ state in the absence of nanodisc. Similar observations on the lack of strong interactions between αSN and DMPC nanodiscs have been made previously (87, 89). One interesting observation is that substantial uptake differences were not observed for the αSN/DMPC complex even at the shortest (millisecond) HDX labeling times (Figure S3). This could imply that the turnover rate of the complex is very fast (*i.e*., with a relaxation time on the order of 10 ms or less) and favors the unbound state, but it could also indicate that complexation with DMPC (which is supported by the SEC analysis) has little impact on the αSN conformational ensemble.

Next, the αSN/(POPG:DMPC (1:1)) nanodisc complex was examined. The POPG lipid content introduces more negative charge on the surface of the nanodiscs, which should provide a more favorable charge environment for αSN docking (87, 89). The inclusion of 50% POPG in the nanodiscs resulted in more prominent changes in αSN deuterium uptake, corresponding to a more compact and/or restricted conformational ensemble (Figure 4, middle) (43). Of the regions detected, the strongest impact was observed in segment 19 – 28, corresponding to the N-terminal helix reported in the structure of αSN reported by Ulmer *et. al*. (PDB 1XQ8) (35). The region from residues 46 – 57 also appears to be weakly, but significantly impacted. This region contains the N-terminus of helix II, which is thought to play a key role in the formation of off-pathway amyloidogenic intermediates of αSN. Decreased uptake in this region could correspond to stabilization and extension of the NAC helix, which has been proposed to occur in αSN-membrane interactions (36, 90, 91).

For the αSN/pure POPG nanodisc complex, substantial decreases in uptake were observed, mainly toward the C-terminus of helix I and the N-terminus of helix II with no changes in the inter-helix loop (covered by segment 39 – 43 in our data). This suggests that the region bounded by the C-terminal end of helix I and the N-terminal end of helix II undergo a substantial change in dynamics, corresponding to a local tightening of the conformational ensemble, likely due to stabilization of helical structure, upon interaction with negatively charged membranes. It is important to note that the C-terminal tail showed no significant variation in deuterium uptake upon binding in all three data sets (Figure 4) which is consistent with most accepted models for αSN aggregation (3, 4, 21, 36, 92).

## Discussion

The consensus view that amyloidogenesis arises from a process of nucleated polymerization driven initially by changes in the secondary structure of the protein has been generally accepted for more than 20 years (92, 93). In the case of αSN, the multitude of conformational modes accessible through interactions with biological membranes has proven of particular importance to the pathology of PD and other synucleinopathies (44, 89, 91, 94). However, this has made the molecular mechanisms of the protein-lipid interaction and the origins of neurotoxicity hard to pinpoint, since slightly different experimental conditions, like lipid composition, lipid-to-protein ratio and type of membrane mimetic greatly vary experimental results (44, 95).

Non-mechanistic studies have demonstrated that the aggregation propensity of αSN correlates with the affinity of the monomeric form of the protein to particular lipids. For example, lipids with neutral or positively-charged head-groups do not influence the rate of amyloidogenesis, but the presence of anionic phospholipids was found to initiate, accelerate - or in some cases inhibit - αSN aggregation (44, 96). These results are supported by the model that arises from our mechanistic data (Figure 6), which indicate that, while αSN may interact transiently with neutral headgroup membranes, these interactions have no impact on its conformational ensemble (Figure 6a).

Computationally-derived mechanistic models have proposed that lipid-protein complexation is driven initially by electrostatic interactions between basic residues and the negatively charged bilayer as well as well as enthalpic gains from increased intrachain hydrogen bonding associated with stabilized helicity (89, 97). One thing on which these models tend to agree is that αSN binding to anionic lipid bilayers is mediated mainly by the first 90 amino acids of the sequence, in an interaction that does not involve deep penetration of the peptide chain below the phospholipid membrane surface (89).

The ambiguity in knowing what increases or decreases amyloidogenic propensity of αSN upon lipid interactions highlights the importance of acquiring a detailed, molecular understanding of how αSN interacts with different lipids. Our work broadly supports the common features of the most widely accepted models for αSN/lipid membrane interactions, including the requirement for negatively-charged head groups to facilitate strong (or, from our view ‘low turnover’) interactions, and the impacts on conformation and dynamics being felt mainly in the N-terminus. However, we are also able to provide a more detailed picture of the interaction that gives direct experimental support for a subset of speculative / computational models (31, 36, 98).

Our millisecond HDX data highlight the N-terminal region as being the most strongly impacted by lipid membrane interactions, which is in agreement with several NMR studies (3, 31, 88, 99). More specifically, we note a substantial change in uptake in the segment 19 - 28 (AEKTKQGVAE), which is in a region proposed to form a stable helix in the initial αSN/membrane interaction This interaction is thought to occur in both ‘physiological’ and pathological models for αSN interactions with synaptic vesicles. One differentiating feature of the ‘physiological’ and ‘pathogenic’ membrane interaction model is that the former involves stabilization of the interhelix loop (LYVGSKTK), amino acids 37-45, to form an ‘extended helix’ structure. This structure preferentially dissociates from the membrane to form weakly amyloidogenic ‘free’ αSN instead of undergoing self-association into pre-fibrillar aggregates. Interestingly, we observe no change in the 39 – 43 segment, which would be included in the ‘extended’ helix under the physiological model. Our data are therefore not consistent with the occurrence of significant ‘helical extension’ in the N-terminal region.

At the same time, we do detect significant ‘tightening’ of the conformational ensemble, consistent with increased helicity, in the 45 – 57 region when the nanodisc membrane model includes POPG (and more strongly when the membrane model is 100% POPG). This provides experimental support for the occurrence of the proposed ‘broken helix’ state, which involves one membrane-bound helix I at residues 1 to 37 and a second, free helix II at amino acids 45-95. It also supports the notion that the broken-helix state is the most populated conformational ensemble when αSN binds to ‘target’ membranes, reinforcing the idea that transitions leading down the amyloidogenic pathway, which are proposed to require melting of helix II, are rare events (Figure 5b and c).

**Figure 5.**
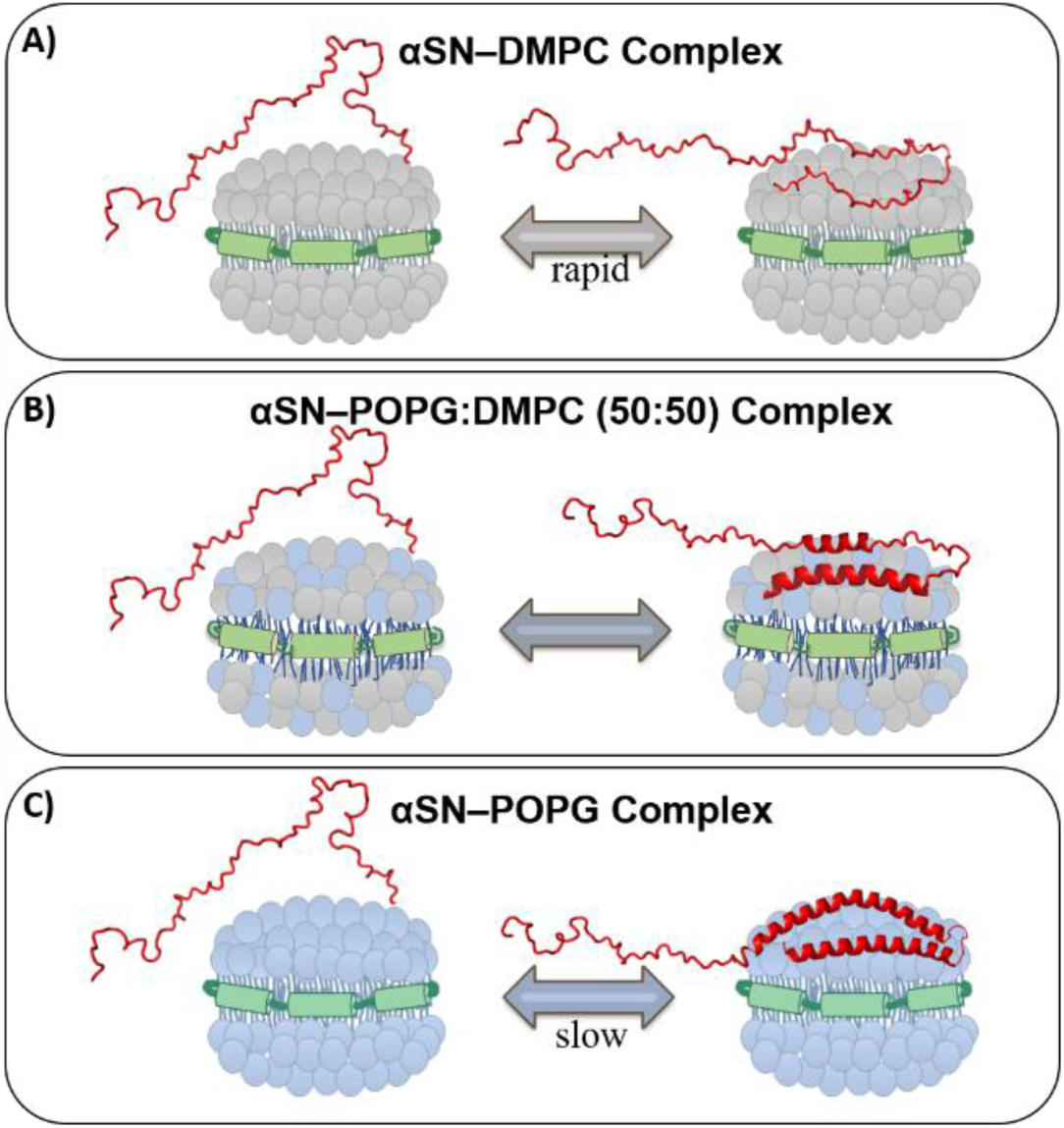
Cartoon representation of the αSN with the corresponding lipid membrane bilayer: A) DMPC nanodisc, B) POPG:DMPC (1:1) nanodisc, and C) POPG nanodisc. αSN represented by PyMol structure (red), POPG lipid by blue circles, DMPC lipids by grey circles and MSP1D1 ΔH5 belt by green cylinders.

Finally, our data are entirely consistent with the widely held view that the C-terminal region, spanning from residue 90 to 140, is largely unaffected by (and uninvolved in) membrane interactions and remains, as in the ‘free’ protein, largely disordered (14, 20).

## Conclusions

With an estimation of nearly nine million people being affected by PD by the year 2030 (95, 100), the necessity to grasp the mechanism(s) of αSN’s oligomerization as well as its core biological function(s) is increasingly vital (93). In this work, we have demonstrated that TRESI-HDX-MS can provide unique insights into conformational dynamics in specific regions of αSN upon complexation with nanodisc membrane mimetics. Specifically, we acquired direct experimental evidence supporting most features of the most widely accepted model of αSN/membrane interactions, particularly direct interactions with the N-terminal region, the occurrence of a dominant ‘broken helix’ conformational ensemble and a lack of involvement for the C-terminus. We further demonstrated that the population of the ‘broken helix’ ensemble is linked to the extent of negative charge in the ‘target’ membrane and that neutral membranes have no significant impact on the αSN conformational ensemble, despite evidence for transient interactions.

Given the relatively straightforward nature and wide applicability of TRESI-HDX-MS studies with nanodiscs, the current work provides a foundation to investigate a broad range of αSN and membrane variants with important phenotypes. For instance, it would be of a great interest to probe conformational dynamics of lipid-protein interactions in αSN mutations associated with autosomal dominant PD (*e.g*. A18T, A29S, A30P, E46K, H50Q, G51D, A53T and A53E). Post-translational modifications, like acetylation and phosphorylation, generate another level of complexity in the structural biases associated with αSN’s amyloidogenic membrane interactions and can easily be explored using the methods introduced here.

## Experimental procedures

### Protein purification

αSN was expressed and purified from *E. coli* BL21 cells containing pET-23a vector encoding the *SNCA* gene. Purification was done by boiling the post-lysis cell pellet in 100 °C bath for 15 min followed by a step gradient anion exchange chromatography with NaCl - 100 mM, 150 mM, 200 mM, 250 mM, 300 mM and 500mM. αSN was concentrated and dialyzed into 20mM Tris-Cl, pH 7.4 using a 3kDa Vivaspin column (Millipore Sigma™ Amicon™). Protein was stored in 20% glycerol at −80°C prior to MS analysis or formation of αSN-nanodisc complex.

MSP1D1 ΔH5 was expressed and purified from *E. coli* BL21-CodonPlus (DE3) cells containing pET28a-MSP1D1delltaH5 vector (Addgene cat# 71714 was kindly purchased by Dr. Audette Lab). In short, purification was performed by running post-lysis supernatant through Ni^2+^–affinity gravity chromatography with three consequent washes of 250 mL of 40 mM Tris/HCl, 0.3 M NaCl, 1% Triton X-100, pH 8.0, 250 mL of 40 mM Tris/HCl, 0.3 M NaCl, 50 mM Na-cholate, 20 mM imidazole, pH 8.0, and 250 mL of 40 mM Tris/HCl, 0.3 M NaCl, 50 mM imidazole, pH 8.0. Elution of MSP1D1 ΔH5 was completed by running 150 mL of 40 mM Tris/HCl, 0.3 M NaCl, 0.4 M imidazole, pH 7.0 through the column. MSP1D1 ΔH5 was concentrated and dialyzed into 20 mM Tris/HCL, 0.1 M NaCl, 0.5 mM EDTA, pH 7.4 using a 10 kDa Vivaspin column (Millipore Sigma™ Amicon™). Protein was stored in 20% glycerol at −80°C prior to nanodisc formation.

### Nanodisc formation

Nanodiscs were made according to established protocols (53, 101). This study involved three types of lipid constituent for nanodisc: 100% 1,2-dimyristoyl-snglycero-3-phosphocholine (DMPC), 100% 1-palmitoyl-2-oleoyl-sn-glycero-3-phospho-(1’-rac-glycerol) (POPG) and 50:50 DMPC:POPG. All three types of nanodiscs were assembled using MSP1D1 ΔH5 as a belt. In short, nanodisc assembly was performed by resuspending each lipid (DMPC and POPG) or lipid mix (DMPC:POPG 50:50) with HPLC grade water for a final concentration of 100 mM usually referred as lipid stock. Then each lipid stock was mixed with 300mM Na-cholate in 1:1 v/v ratio and heated under ∼50° tap water for 2-3 minutes, as well as, sonicated in an ultrasonic bath until the solution was clear and no lipid remained on the walls of the tube. The scaffold-to-lipid molar ratio was calculated from geometrical considerations. Upon mixing the calculated ratios of DMPC, POPG and DMPC:POPG (50:50) lipids with MSP1D1 ΔH5 scaffold protein, these nanodisc reconstruction mixtures were incubated for 15 minutes at 25°C, then 1 hour at 4°C. Next, BioBeads SM-2 (Bio-Rad Laboratories) in the amount of 0.6 g per 1 mL were added to the reconstruction mixture and the resulted mixes were incubated for 2 hours at 4°C for DMPC nanodiscs and 4 hours at 4°C for POPG and DMPC:POPG (50:50) nanodiscs on the orbital shaker. Formed nanodiscs were then removed from the beads and stored at 4°C for future applications. The formation of nanodiscs was confirmed by size-exclusion chromatography (SEC). All lipid were purchased from Avanti Polar Lipids, Inc.

### αSN - nanodisc complex assembly

The assembly of the complex, αSN with the DMPC or POPG or DMPC:POPG (50:50) nanodiscs (MSP1D1 ΔH5 belt for all), was made by mixing nanodisc with the protein in a 1:1 molar ratio and incubating this mixture at 37°C for 1 hour at 100 rpm. Formed complex was verified and purified from unbound lipids and proteins by SEC using Tosoh Biosciences TSKgel BioAssist G3SWXL 7.8 × 300 mm (5 μm, 250 Å) column. Fraction were eluted with 100 mM ammonium acetate, pH 7.0 at 0.5 mL/min and further analyzed by SDS-PAGE. Assembled complex was stored on for further MS applications.

### TRESI-HDX-MS experiments

TRESI-HDX-MS experiments were completed according to the experimental setup in Figure 1 and established protocols (59, 65, 102). Flow rate parameters were: 5 µL/min for 10 µM αSN alone or 10 µM complex, 5 µL/min for D_2_O, and 15 µL/min for 10% acetic acid. Quenched samples were digested by protease XIII, and the resulting peptides were ionized by electrospray into the Synapt G2S mass spectrometer (Waters, MA). IMS was employed in the TriWave cell to improve spatial resolution of peptides in the digested sample. Six time points (0.12 sec, 0.34 sec, 0.66 sec, 1.32 sec, 2.2 sec and 3.3 sec) were obtained for each protein state – free αSN and the complex of αSN with three different lipid constituents. MS parameters for TRESI-HDX-MS runs were: 2.5 kV capillary voltage, 120°C source temperature, 25.0 sampling cone, 250 °C desolvation temperature, 50 L/hr cone gas flow, 6.0 Trap collision energy (CE), 6.0 Transfer CE and 90 mL/min IMS gas flow. Acquisition range was m/z 400 to 1500. TOF analyzer was in positive sensitivity mode.

### HDX data analysis

Identified peptides were analyzed for deuterium uptake using Mass Spec Studio 1.0 - a free software developed in Dr. David Schriemer’s research group (103). Deuterium uptake difference and kinetic plots were made using Excel software. Mapping of the deuterium uptake onto crystal structures of αSN was performed using PyMOL software (104).

## Acknowledgements

We would like to acknowledge Dr. Gerald Audette at York University for kindly providing plasmids for MSPs.

## Funding and additional information

This work was supported by the Natural Sciences and Engineering Council of Canada Discovery (DG-544760) and Collaborative Research and Development (CRDPJ-480432) grant programs.

